# Constitutive and variable patterns of genome-wide DNA methylation in populations from spatial-environmental range extremes of the bumble bee *Bombus vosnesenskii*

**DOI:** 10.1101/2023.05.02.539175

**Authors:** Sarthok Rasique Rahman, Jeffrey D. Lozier

## Abstract

Unraveling molecular mechanisms of adaptation to complex environments is crucial to understanding tolerance of abiotic pressures and responses to climatic change. Epigenetic variation is increasingly recognized as a mechanism that can facilitate rapid responses to changing environmental cues. To investigate variation in genetic and epigenetic diversity at spatial and thermal extremes, we use whole genome and methylome sequencing to generate a high-resolution map of DNA methylation in the bumble bee *Bombus vosnesenskii*. We sample two populations representing spatial and environmental range extremes (a warm southern low-elevation site and a cold northern high-elevation site) previously shown to exhibit differences in thermal tolerance and determine positions in the genome that are constitutively and variably methylated across samples. Bisulfite sequencing reveals methylation characteristics similar to other arthropods, with low global CpG methylation but high methylation concentrated in gene bodies and in genome regions with low nucleotide diversity. Differentially methylated sites (n = 2,066) were largely hypomethylated in the northern high-elevation population but not related to local sequence differentiation. The concentration of methylated and differentially methylated sites in exons and putative promoter regions suggests a possible role in gene regulation, and this high-resolution analysis of intraspecific epigenetic variation in wild *Bombus* suggests that the function of methylation in niche adaptation would be worth further investigation.

## Introduction

Understanding the ecological and evolutionary mechanisms of adaptation to complex ecological niches is a central goal of evolutionary genomics^1–3^. Species with large geographic distributions face diverse pressures from environmental heterogeneity across populations^4, 5^, and genotypic and phenotypic variation among dissimilar environments can provide the raw material for local adaptation^6^. Species in mountainous regions, in particular, can experience extreme variations in abiotic conditions such as temperature, precipitation, or air density^2, 7, 8^. Population-level genomic changes at the spatial-environmental extremes in widespread montane species could thus improve our current understanding of how species tolerate diverse bioclimatic conditions and provide insights into potential mechanisms of adaptability and robustness under global climate change^6, 9, 10^.

DNA sequence-based variation has been the most commonly examined form of genomic adaptation in wild populations, however, epigenetic variation, such as DNA methylation, histone modifications, and regulatory non-coding RNAs, is increasingly recognized as a potential mechanism of rapid environmental adaptation or plasticity^11–13^. Epigenetic mechanisms can generate flexible responses to various environmental stimuli without modifying genome sequences, and they are potentially important for species that occupy diverse bioclimatic niches^14^. Cytosine (CpG context) methylation is the most prevalent form of epigenetic methylation^15^, however, the extent of CpG methylation and its functional significance varies substantially across lineages^16^. For example, mammals exhibit higher (70-80%) of global CpG methylation^17^ compared to plants (4-40%)^18^ and arthropods (<1% to 14%)^19^. While in plants, DNA methylation primarily occurs in repetitive regions, especially in transposon elements (TEs)^20^, in mammalian genomes, cytosine methylation is consistent except in the CpG islands (i.e., CG motif-rich genomic regions) near promoters and transcription start sites (TSS)^21^. Mammalian CpG methylation has been linked to various molecular functions^17^, such as gene silencing, genomic imprinting, and stabilization of regulation of gene expression ^22–25^. In arthropods, methylation functionality has been attributed to varied biological processes such as reproduction^26^, caste determination^27–29^, and regulation of gene expression via differential exon usage^30^. Arthropod CpG methylation is most prominent in gene bodies compared to intrageneric and intergenic regions, but levels vary widely across lineages^31^. For example, model organism *Drosophila melanogaster* has very low amount of CpG methylation^19, 32^ due to the absence of a key methyltransferase gene (*Dnmt3*)^33^. Characterizing genome-wide patterns of DNA methylation across a wide range of taxa^34^ will be important in understanding the distribution of constitutive CpG methylation patterns across multiple lineages and identifying the extent of intraspecific epigenomic variability. The function of such variable epigenetic changes may be especially relevant in the context of adaptation to anthropogenic climate change.

Bumble bees are among the most economically and ecologically important pollinating insects^35, 36^ that primarily inhabit cool temperate, alpine, and arctic climates^37^. Bumble bees exhibit remarkable phenotypic and physiological adaptations for thermal regulation^38^, such as an insulated pile, generating heat through shivering of flight muscles, and shunting mechanisms that prevent overheating^39–41^. Such thermal adaptations allow bumble bees to fly and forage in diverse thermal niches than many other insects^42, 43^. Like many insects^44^, many bumble bee species have declined in geographic range and abundance^45^, seemingly driven at least in part from anthropogenic climate change^46, 47^. In North America, while several bumble bees have recently declined dramatically, many species remain common ^48–50^, and species-specific responses to global climate change^46^ indicate that some species may tolerate warming temperatures better than others^51^. The nature of genomic and epigenomic variation within species that occupy diverse environments will be valuable for understanding why species may be vulnerable or resistant to climate change.

*Bombus vosnesenskii* is a common bumble bee species that is distributed across latitudinal and altitudinal gradients in Western North America, principally in California, Oregon, and Washington, USA (Fig. 1). Population genetic studies have found low levels of intra-specific genetic differentiation and weak population structures across the *B. vosnesenskii* range^5, 52^, and *B. vosnesenskii* is one of two North American bumble bee species projected to expand its range under future climate change scenarios^51^. Therefore, studying environment-associated genomic variation may provide insights into species-specific responses that may mitigate the negative impacts of climate change. As a widely distributed and ecologically crucial native pollinator, *B. vosnesenskii* has gained substantial attention as a research subject for population genetics^52, 53^, pollination biology^54^, and abiotic adaptation^55, 56^ studies. Genome scans across a broad latitudinal and altitudinal range using restriction site-associated DNA sequencing (RAD-Seq) and environmental association analysis identified relatively few potential genomic regions associated with thermal and desiccation tolerance^55^. However, analysis of thermal tolerance across latitude and altitude extremes of the *B. vosnesenskii* range provided some evidence for local adaptation, with population-level variation in lower thermal tolerance (CT_MIN_) of laboratory-reared bees that matched the annual temperature of respective source populations^56^. Moreover, transcriptional differences among populations were detected at these lower CT_MIN_ thresholds. In contrast, there was no evidence of intrapopulation variation in responses to upper thermal limit (CT_MAX_), suggesting evolutionary conservation of physiological and molecular responses under heat stress^56^. Results from these studies provide a foundation for investigating other types of variation that may contribute to molecular responses, including epigenetics, which will contribute to a greater understanding of potentially adaptive thermal tolerance mechanisms in this species.

**Figure 1.**
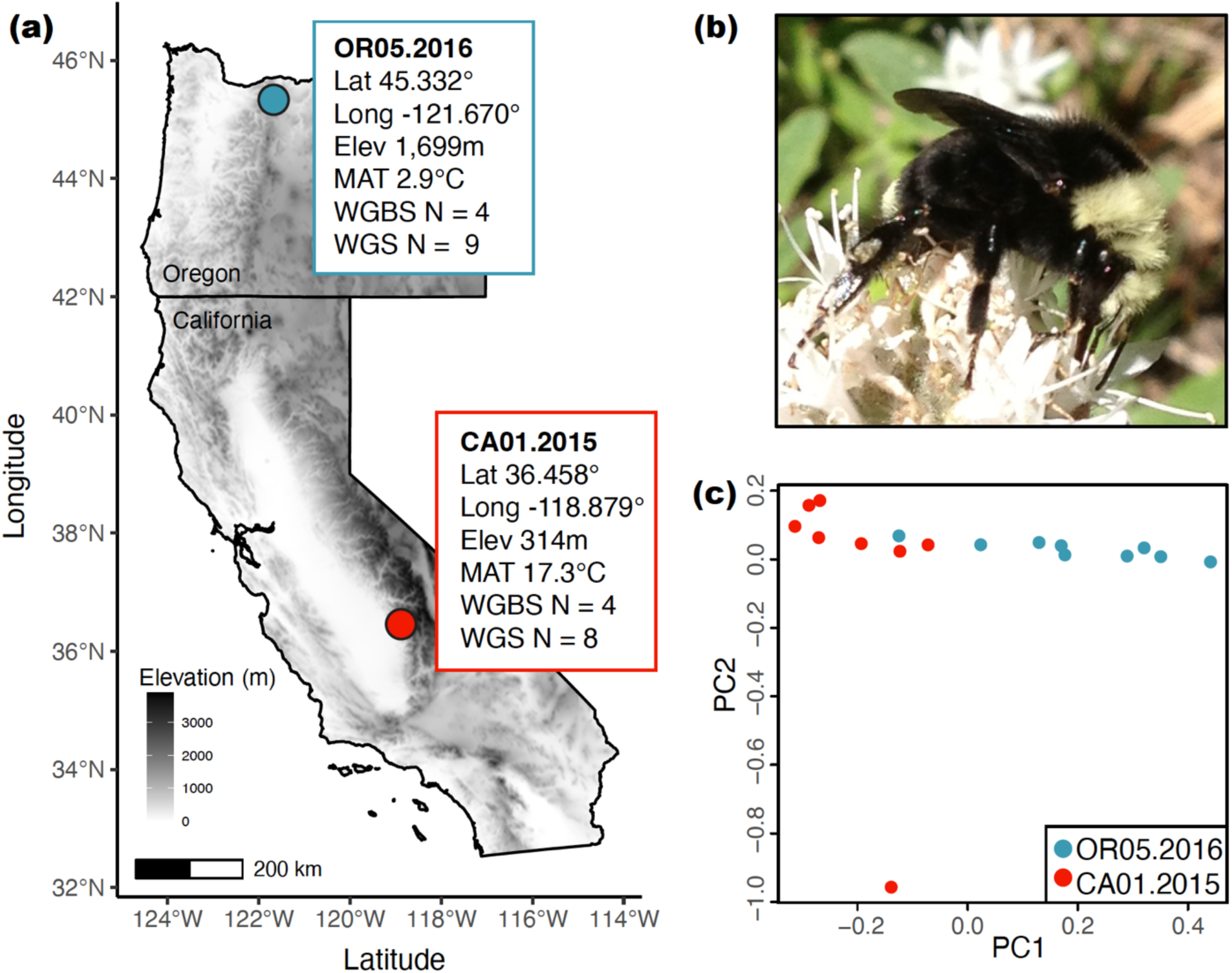
**(a)** Map, spatial information (latitude, longitude, elevation, and mean annual temperature from the WorldClim v.2^135^), and sample sizes for bisulfite (WGBS) and whole genome (WGS) sequencing for the two *B. vosnesenskii* study populations. **(b)** Photograph of *B. vosnesenskii*, **(c)** Genome-wide Principal Components Analysis (PCA) from covariances estimated by PCAngsd for the *B. vosnesenskii* populations sampled for WGBS.

The majority of epigenetics and DNA methylation studies in bumble bees have centered around determining its role in sex/caste determination^57, 58^, reproduction^59^ and development^60^ using lab-reared individuals of two commonly used model species, *B. terrestris*, and *B. impatiens*^61^. However, little is known regarding the role of epigenetics in shaping niche-specific thermal adaptation in wild bumble bees, which might provide insights into the adaptive variation that could allow responses to environmental variation^2, 62, 63^. The availability of reference genomes for multiple bumble bee species^64, 65^ now facilitates expanding the phylogenetic scope of methylation research in bumble bees. In this study, we use very high-coverage whole genome bisulfite sequencing (WGBS) data to map epigenetic variation in *B. vosnesenskii.* We also evaluate the potential for intraspecific epigenomic variation by sequencing populations representing the spatial and thermal range extremes, focusing on wild-caught samples taken from two extreme populations: a southern low elevation population from California, USA (warm extreme) and a northern high elevation population from Oregon, USA (cold extreme) (Fig.1). In addition to detailed characterization of the methylome of the species overall and testing for intraspecific epigenetic differentiation, we also assess possible relationships between methylation with population genetic diversity or structure using single nucleotide polymorphisms (SNPs) from whole genome sequencing (WGS). Specifically, we aim to: (i) characterize the major trends in constitutive methylation patterns in *B. vosnesenskii* and identify putative major functions related to genome-wide CpG methylation; (ii) compare and contrast epigenetic profiles from populations at latitude and altitude extremes to assess variability in the methylome and characterize the genomic location and potential functional roles of differentially methylated CpGs; and (iii) investigate the potential relationship between population genetic diversity and genome-wide CpG methylation levels in *B. vosnesenskii*. Our research provides insights into the distinct nature of constitutive and variable DNA methylation in populations from the spatial-environmental range of *B. vosnesenskii*, and it also highlights the existence of intraspecific epigenetic variation that may aid in generating regional variation in genotypes and phenotypes to adapt the species across a range of intricate biological niches.

## Results

### Constitutive patterns CpG methylation across *B. vosnesenskii* genome

Overall CpG methylation across the genome was 1.1% ± 0.9% SD which was calculated from the percent methylation per CpG cytosine values across all samples (Fig. 2a). The low-elevation California (CA) population had slightly higher percent methylation (1.17% ±0.06% SD) than the high-elevation Oregon (OR) population (1.03% ± 0.04% SD) (Fig. 2a). Most sequenced CpGs (methylated + unmethylated) were located in introns (57.90%) and intergenic (23.42%) regions, with 5.73% in coding sequences (CDS) and 5.09% in untranslated regions of exons (exon UTRs) (Fig. 2b). The distribution of methylated CpGs varied substantially from the overall distribution of CpGs, with both highly methylated (>50% average methylation; n = 112,996, ∼0.78% of all CpGs) and sparsely methylated (10-50% methylation, n = 186,846, ∼1.28% of all CpGs) sites predominantly found in CDS (Fig. 2b). Specifically, 70.85% of sites that were classified as highly methylated in all samples were in CDS, 13.02% were in introns, 9.50% in exon UTRs, and much lower percentages in the remaining annotation feature categories (0.76-3.13%) (Fig. 2b). Although highly methylated CpGs are only ∼0.78% of all CpG positions in the genome, the proportion of highly methylated CpGs per total sequenced CpGs in CDS is even more extreme (9.36% of all CpGs in CDS) compared to introns (0.17%) and intergenic (0.04%) regions. Annotation feature-specific distributions of highly methylated CpGs is significantly different from distribution of all CpGs (Pearson’s Chi-squared test with Yates’ continuity correction; FDR corrected P < 0.05) for all eight annotation features [i.e., exon UTR, CDS, intron, upstream flank, downstream Flank, long non-coding RNA, transposable elements (TE), intergenic].

**Figure 2.**
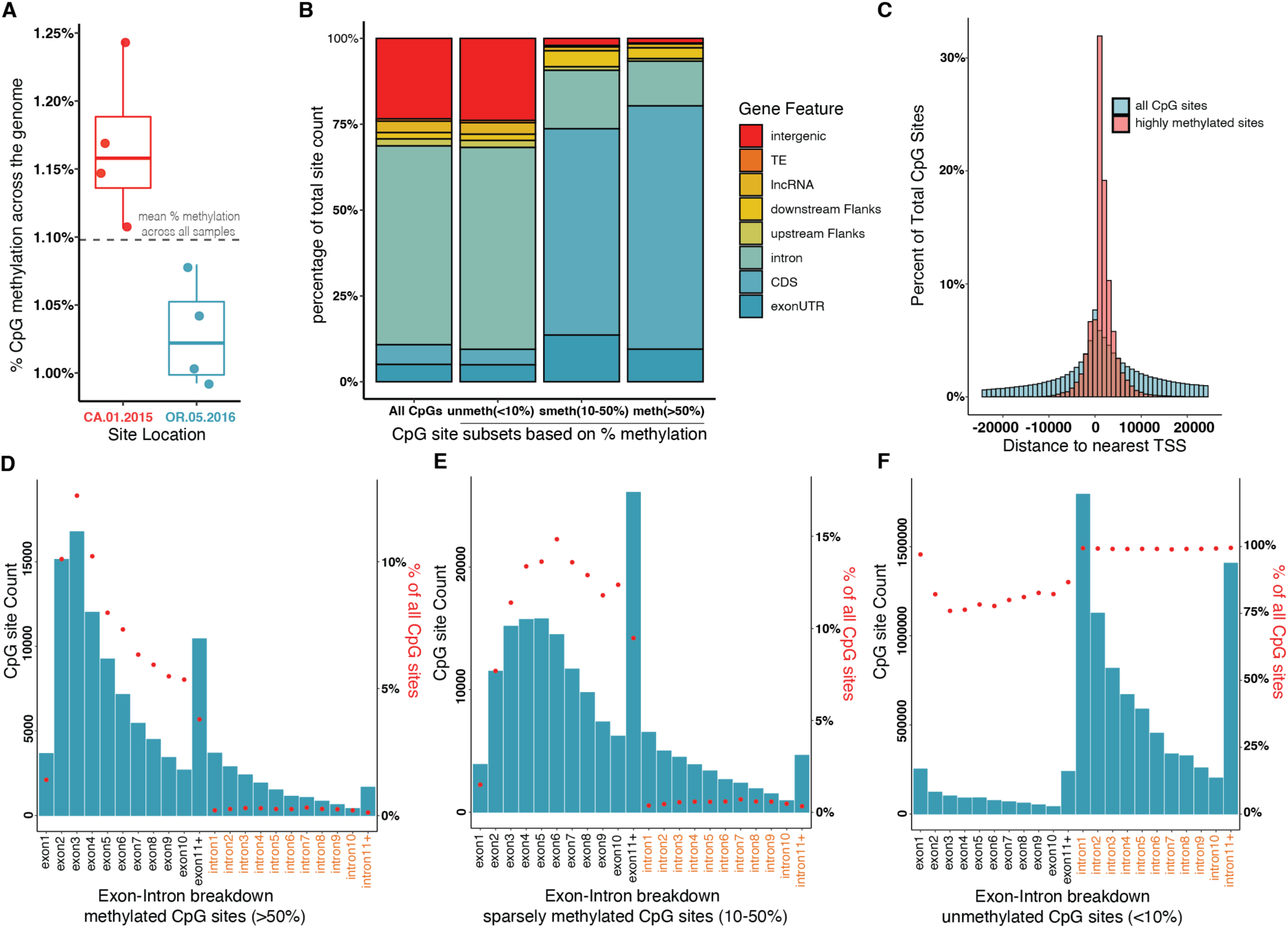
General Pattern of genome-wide methylation in *B. vosnesenskii* study samples. **(a)** Box plot exhibiting sample specific per base percent methylation for the CpGs present in every sample. **(b)** Bar plots of genomic feature-based annotation proportions for all CpGs, unmethylated sites, sparsely methylated sites, and highly methylated sites. **(c)** Histogram of distances to nearest TSS for all CpGs and highly methylated sites. d-f. Exon intron breakdown of gene body methylation for highly methylated **(d)**, sparsely methylated **(e)**, and unmethylated CpGs **(f)**. Y-axis blue bars represent the actual count, and red dots depict the proportion of individual genomic features (i.e., exons and introns) relative to similar annotation feature counts for all CpGs.

Consistent with the overabundance of methylated sites in CDS, a greater number of highly methylated sites clustered near the transcription start site (TSS) than predicted from the genome-wide distribution of TSS distances for all CpGs (Fig. 2c), with the absolute mean distance from TSS for highly methylated CpGs was much shorter (2,438.78bp) compared to the absolute mean distance from TSS for all CpGs (27,981.11bp). There were ∼4.5 times more CpGs in downstream (gene bodies and 5′ UTR) of TSS (n = 92,403) compared to the number of CpGs in upstream (e.g., likely promoter regions) of TSS (n = 20,561), which is substantially higher than for all CpGs [∼1.51x more sites in downstream of TSS (n = 8,708,196) compared to the CpGs in upstream (n = 5,774,550)]. The distribution of distances to the TSS for highly methylated sites was significantly different than that for all CpGs (two-sided two-sample Kolmogorov-Smirnov test, D = 0.35738, P < 2.2e-16). The distribution of sparsely methylated CpGs was similar to that of highly methylated CpGs (Fig. 2b). As expected, unmethylated CpGs (<10% methylation average methylation; n = 14,283,650, ∼97.94% of all CpGs) largely matched that of the genome-wide distribution of CpGs except for a slightly smaller proportion in CDS (due to the greater methylation presence in CDS).

To examine the distribution of methylation levels relative to CpG background within genes, we examined the frequencies of highly methylated, sparsely methylated, and unmethylated CpGs for exons, introns and other annotation features (Fig. 2d-f). The first clear pattern is that exons have much greater levels of methylation, both in absolute numbers of methylated CpGs and even more clearly apparent when visualized as a percentage of available CpGs per feature (Fig. 2d-e). For exons, exon 2-4 harbored substantially more highly methylated sites (10.1%, 12.6% and 10.2% relative percentages compared to all CpGs, respectively) than the first exon (1.4%), and generally decreased from exon 3 through the remaining exons (Fig. 2d). A similar pattern was apparent in the sparsely methylated sites, although the distribution was not as sharply biased toward exons 2 and 3 (Fig. 2e). In contrast, the exon-specific distribution pattern is reversed in unmethylated sites (Fig. 2f), as the first exon has more unmethylated sites (97%) than exon 2 (82%), exon 3 (76%) or rest of the exons, although as discussed above the number and proportion of unmethylated CpGs is reduced in exons relative to introns overall. For introns, there was a downward trend in raw counts from upstream to downstream intron locations across the gene body for all three (methylated, sparsely methylated and unmethylated) categories, however, the trend is absent when considered as percentages of available CpGs (Fig. 2d-f). We separately evaluated patterns in long non-coding RNAs, which showed a similar exon-intron breakdown.

We also visualized the chromosomal distribution of CpGs across the genome (Fig. 3). Most of the CpGs across the genome have low methylation (<10%) and highly methylated sites are relatively scarce (Fig. 3a), however, plotting the average per base percent methylation across the genome shows their distribution is non-random as we discovered the large regions of very low methylation in genomic scaffolds punctuated with peaks of methylation heavy regions (Fig. 3c-d); gene-specific visualization of CpGs (Fig. 3e-f) exhibits that this distinct pattern of clustering of highly methylated CpGs are predominantly located in gene bodies.

**Figure 3.**
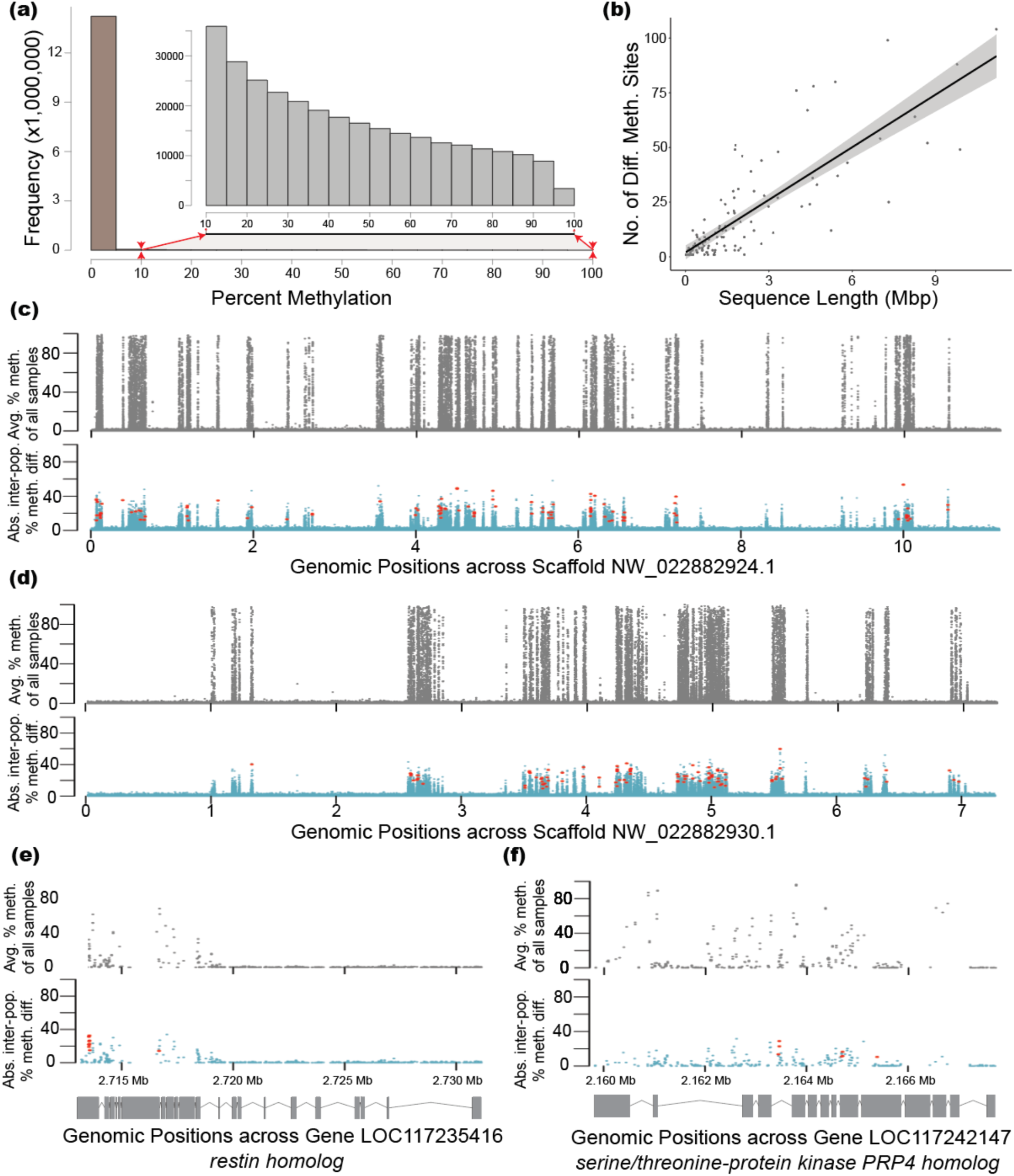
Genome-wide distribution of CpG methylation in *B. vosnesenskii*. **(a)** Frequency histogram of percent methylation of all CpGs with the distribution of CpGs with more than 10% methylation zoomed-in inset plot. **(b)** Scatter plot of correlation between scaffold length (Mbp) and the number of differentially methylated sited harboured in the individual scaffolds. **(c-f)** Manhattan plots of the genomic landscape of average percent methylation (top panel) across all samples and absolute inter-population percent methylation difference (bottom panel) across scaffold NW_022882924.1 **(c)**, scaffold NW_022882930.1 **(d)**, *restin homolog* **(e)** and *serine/threonine-protein kinase PRP4 homolog* **(f)** with their exon-intron gene structures.

### Patterns of differentially methylated CpGs between populations from spatial-environmental range extremes of *B. vosnesenskii*

The principal component analysis (PCA) of all methylated CpGs showed that 31.44% of the variation was explained by first two principal components with weak separation of OR and CA samples, and greater variation within CA, although population-specific clustering was more prominent in a clustering dendrogram (When analyses were repeated using CpGs that were variably methylated among all samples (excluding sites within 2 SD of average per base percent methylation, n = 901,868 CpGs) there was much clearer evidence of population-specific clustering (Fig. 4a), and hierarchical clustering also exhibited distinct population-specific clusters (Fig. 4b). PCA and hierarchical clustering analysis using only differentially methylated sites, obviously indicated clear distinction between two populations.

**Figure 4.**
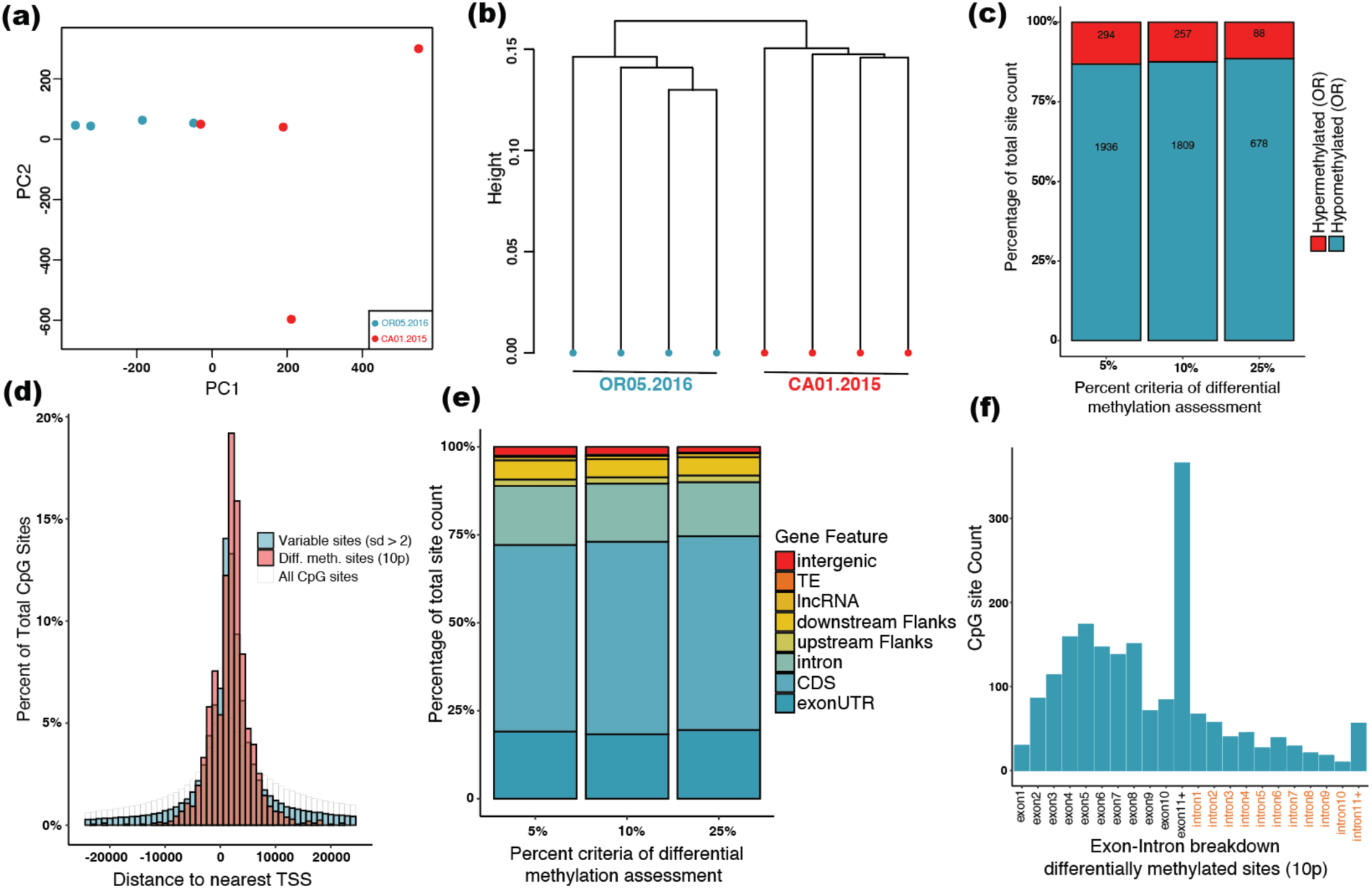
General patterns of clustering and distribution of variable (SD >2) and differentially methylated CpGs in *B. vosnesenskii*. **(a)** Principal Component Analysis (PCA) of methylation profiles of variable CpGs (SD >2). **(b)** Hierarchical clustering of methylation patterns of variable CpGs using Ward.D2 algorithm **(c)** Bar plot of counts and percentages of hypermethylated and hypermethylated CpGs in high elevation (Oregon) samples assessed using three different criteria (5%, 10%, and 25% methylation difference). **(d)** Histogram of distances to nearest TSS for variable (SD >2), differentially methylated, and all CpGs. **(e)** Genomic feature-based bar plots depicting annotation proportions for differentially methylated sites assessed at three criteria (5%, 10%, and 25% methylation difference). **(f)** Exon-intron breakdown of gene body methylation for differentially methylated CpGs.

We identified 2,066 significantly differentially methylated sites (≥10% methylation difference, FDR corrected q ≤ 0.01) between OR and CA. Most (n= 1,809; 87.6%) were hypomethylated in OR relative to CA. Differential methylation at 5% and 25% difference thresholds provided the similar percentages of hypomethylated (86.8 and 88.5%, respectively) sites in OR (Fig. 4c). This result is consistent with the sample-specific methylation frequencies (Fig. 2a) that shows slightly lower overall percent average methylation across the genome in OR.

There was a significant positive correlation between the number of differentially methylated sites and the sequence length of the scaffolds (Pearson’s r = 0.82, P < 0.001; Spearman’s rho = 0.8, P < 0.001)(Fig. 3b), however, as for constitutively methylated CpGs, the distribution within scaffolds was clearly not random (Fig. 3c-f). Differentially methylated sites were distributed even more closely to the TSS (mean = 3622.9bp) than all CpGs (27,981.1bp, two-sided Two-sample Kolmogorov-Smirnov test, D = 0.329, P < 2.2 x 10^-16^) or variably methylated CpGs (absolute mean distance 16304.33bp; two-sided Two-sample Kolmogorov-Smirnov test, D = 0.174, P < 2.2 x 10^-16^) (Fig. 4d). Similar to constitutively methylated CpGs, differentially methylated CpGs are also more numerous downstream of the TSS (n=1,540) than upstream (n= 524), indicating greater abundance in gene bodies compared to the promoters. Differentially methylated CpGs (10% methylation difference threshold) were mostly found in coding sequences (54.72%) and exon UTRs (18.32%) while relatively few were in introns (16.53%) and intergenic regions (2.22%). Similar patterns were exhibited for differentially methylated sites at 5% and 25% thresholds (Fig. 4e). Again, the first exon had fewer differentially methylated CpGs compared to downstream exons (Fig. 4f), and differentially methylated CpGs declined from upstream to downstream across the gene body (Fig. 4f). Long non-coding RNAs also showed more differentially methylated CpGs in exons (n = 92) compared to introns (n = 50). Annotation-specific distributions of differentially methylated CpGs were significantly different from the distributions of all sequenced CpGs (Pearson’s Chi-squared test with Yates’ continuity correction, FDR-corrected P < 0.05) for seven out of the eight annotation features (exon UTR, CDS, intron, downstream Flank, long non-coding RNA, TE, intergenic); only the “upstream” feature was not significant (P = 0.509).

### Genome-wide population structure, genetic diversity and the relationship with CpG methylation

Population structure was weak (F_ST_ = 0.025, 95% CI: 0.025-0.026), but some separation of samples by population was apparent along the first PCA axis (Fig. 1c). Nucleotide diversity (π) per site was similar between the populations, with global π = 0.00197 (95% CI: 0.00196 – 0.00198), OR π = 0.00191 (95% CI: 0.00191 – 0.00192), and CA = 0.00193 (95% CI: 0.00192 – 0.00194), suggesting that differences in genetic diversity do not drive differences in observed methylation levels between populations.

We tested the relationship between general methylation patterns and population genetic diversity across 1kb regions within the *B. vosnesenskii* genome. There was a strong correlation between the mean methylation proportion per CpG per 1kb window (n = 232,788 windows) and both the raw number (Pearson’s r = 0.84, t_232786_ = 737.97, P < 0.001; Spearman’s rho = 0.46, S = 1x#x00D7;10^15^, P < 0.001) and proportion (r = 0.96, t_232786_ = 1642.3, P < 0.001; rho = 0.46, S = 1x#x00D7;10^15^, P < 0.001) of highly methylated CpGs per window. We thus performed analysis only on counts of highly methylated CpGs. The number of highly methylated sites per 1kb window was negatively correlated with π (Fig. 5a) (Pearson’s r = -0.22, t_232786_ = 110.59, P < 0.001; Spearman’s rho = -0.29, S = 2.7x#x00D7;10^15^, P < 0.001). This relationship was not seemingly driven by the number of available CpGs per window, as low diversity windows had fewer CpGs in general (Fig. 5a), so the proportion of CpGs methylated per window thus also declined significantly with π (Fig. 5b; Pearson’s r = -0.24, t_232786_ = -118.88, P < 0.001; Spearman’s rho = -0.29, S = 2.7x#x00D7;10^15^, P < 0.001). There was a weak relationship for F_ST_ and the mean percent methylation difference per CpG per 1kb window between populations (Pearson’s r = 0.01; t_229624_ = 4.97, P < 0.001), although this was not significant for Spearman’s rank correlation (Spearman’s rho = 0.004; P > 0.05) (Fig. 5c). Because above data suggested that certain sites were likely to never be methylated in *B. vosnesenskii* and thus would not differ among populations, it is possible that such regions could affect patterns of differentiation within methylated regions. We thus also evaluated the F_ST_-methylation difference relationships after excluding windows with no methylated CpGs (<10% threshold; n = 22,542 1kb windows retained) and there was no correlation (Pearson’s r = 0.000, Spearman’s rho = 0.000; both P > 0.05; Fig. 5d), suggesting that the weak positive correlation above was likely driven by intragenic or intronic windows with both weak divergence and no methylation.

**Figure 5.**
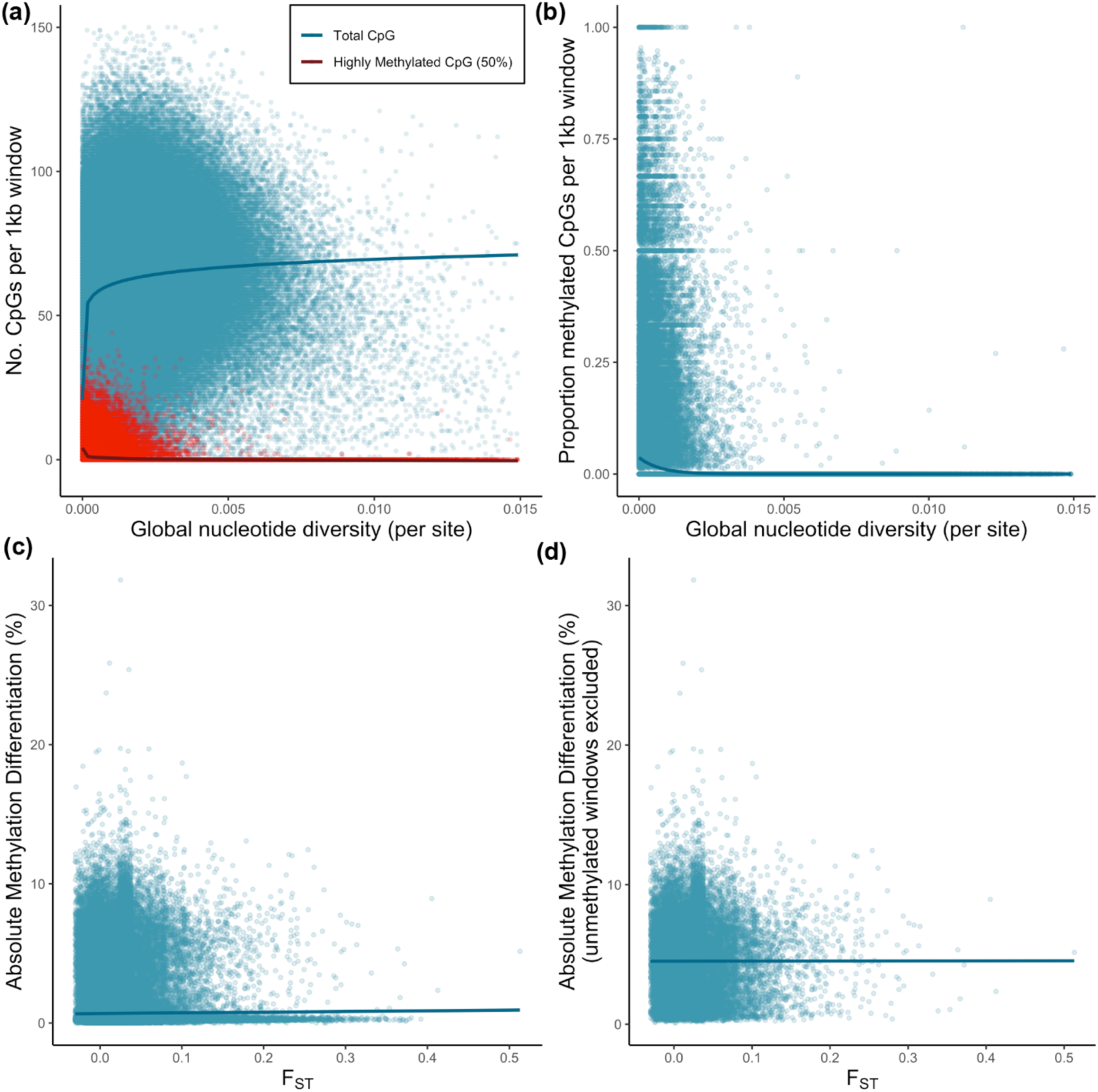
Relationships between genetic diversity from whole genome re-sequencing and methylation patterns. **(a)** Relationship between global nucleotide diversity across samples and the number of methylated CpGs (50% threshold, red) and all sequenced CpGs (blue). Lines fit with a log relationship for visualization. **(b)** Relationship between global nucleotide diversity across samples (log-transformed) and the proportion of methylated CpGs (50% threshold). Line fit with a binomial model for visualization. **(c)** Relationship between F_ST_ and absolute percent methylation differences between populations, **(d)** Relationship between population-level F_ST_ and absolute percent methylation differences excluding 1kb windows with no methylated CpGs. Line fit with a linear relationship for visualization, as no substantial relationship between the variables was detected.

**Figure 6.**
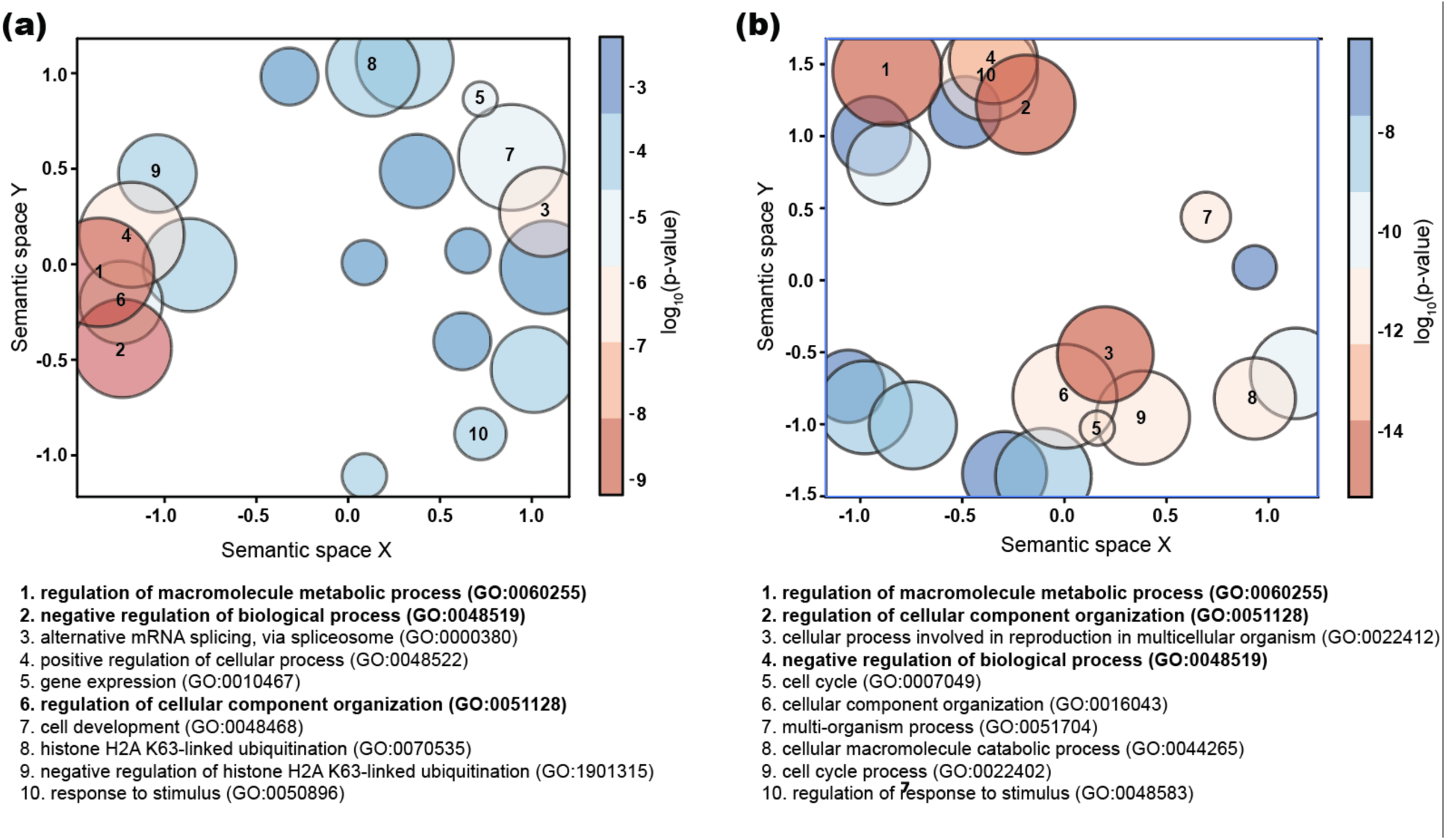
Summarized visualization of biological process-related GO terms from unique genes (n=44) harboring a minimum of 100 highly methylated (>50% methylation) sites (**a**) and unique genes (n=1272) harboring differentially methylated sites assessed at 10% methylation difference (**b**). The top ten biological process-related clusters with their corresponding GO IDs were listed at the bottom of each plot, and common terms between the two lists were boldfaced.

### Gene ontology analysis of genes harboring highly methylated and differentially Methylated CpGs

Analyses of unique genes (n = 44) containing ≥100 highly methylated sites provided 558 statistically significant (FDR corrected q ≤ 0.01) GO terms and a total of 63 summarized GO term clusters. Overall, these GO terms and summarized GO clusters were linked to fundamental cellular activities, such as metabolism, binding, regulation of biological processes, gene expression machinery and cell development. Gene Ontology (GO) analyses of genes (n = 1,272) harboring differentially methylated sites (10% difference threshold) produced 1,473 significantly enriched terms that grouped into 103 clusters that were likewise associated with diverse biological processes, including binding, various enzymatic activities, reproduction, cell cycle, development, metabolism, response to stress and cell communication, and signaling activities. 387 overlapping GO terms from the general and differential methylation enrichment analyses included five overlapping biological process-related clusters [regulation of macromolecule metabolic process (GO:0044265), cell development (GO:0048468), negative regulation of biological process (GO:0048519), regulation of cellular component organization (GO:0051128), regulation of macromolecule metabolic process (GO:0060255)], three molecular function related clusters [chromatin binding(GO:0003682), RNA binding (GO:0003723) and protein binding (GO:0005515)] and six cellular component related clusters. Comparison of GO terms from this study with two previous studies^55, 56^ indicates a functional-level convergence regarding population-specific thermal/environmental adaptation as we noticed several overlapping GO terms, such as, cell signaling and communication, development, reproduction, metabolic functions and response to stimuli, immune function and stress response.

## Discussion

This study presents a high-coverage methylome analysis for the North American bumble bee *B. vosnesenskii* and it is the first to provide initial insights into CpG methylation patterns in wild-caught bumble bees from climatically distinct locations. Genome-wide methylation patterns in *B. vosnesenskii* are similar to those observed in other arthropods and hymenopterans, with a preponderance of highly and sparsely methylated sites found in gene bodies and unmethylated sites disproportionately represented in introns and intragenic regions. We also identified multiple (n=2,066) differentially methylated CpGs between the two sampled populations, predominantly in exons and putative promoter regions, suggesting that epigenetic marks can vary across bumble bee species’ ranges. Our study also reconfirmed previous findings of low genetic diversity and genome-wide genetic homogeneity in *B. vosnesenskii* and showed that while highly methylated regions tended to occur in genome regions with relatively low nucleotide diversity, there was no clear relationship between methylation differentiation and genetic differentiation across genome regions. This in-depth high-coverage analysis of epigenetic variations in *B. vosnesenskii* offers novel biological insights into the factors that may shape the genome-wide distribution of DNA methylation in bumble bees and provides a valuable starting point for more detailed studies of epigenetic mechanisms that may be involved in environmental adaptation or plasticity in this species.

Our first research objective was to characterize the constitutive methylome of *B. vosnesenskii* workers collected from diverse climatic regions within the species’ range to identify features that were highly or rarely methylated in all individuals. The low genome-wide CpG methylation (∼1.1%) is similar to other Hymenoptera^66^, including other bumble bees ^58–60^, the honey bee *Apis mellifera*^67^, the wasp *Nasonia vitripennis*^68^. Such trends are generally common in holometabolous insects^31^ apart from a few unusual instances ^31^. Despite low overall methylation, sparsely distributed peaks of high CpG methylation were non randomly distributed across scaffolds owing to a concentration of methylation in gene bodies, especially exon sequences. This intragenic CpG methylation is also a classic characteristic in insects^19, 28, 67–69,31, 59, 70^ and gene body methylation is likely ancestral^71^. Thus, our results add to the growing body of evidence across the multiple insect orders where the prevalence of gene body methylation was observed irrespective of substantial variability in global methylation levels^31^.

Within genes, methylation was substantially biased towards the 5′ region, with a higher concentration of CpG methylation near the TSS (Fig. 2c) and a relatively gradual decrease of CpG methylation across (5′ to 3′) the transcription unit (Fig. 2d-f). At a more granular level, exon sequences have substantially more methylated sites than introns (Fig. 2d), with a disproportionate distribution of highly methylated sites in exon 2-4, with fewer in exon 1 (Fig. 2d). Similar 5′ biased methylation is observed in bees^67^, wasps^68^, ants^69^ and more generally in holometabolous insects. In contrast, 3′ bias is more prominent in many hemimetabolous insects^72, 73^ and mammals with much higher global methylation^70^. The disproportionate exon-intron breakdown patterns across genes and depleted methylation in the first exon, are also common in Hymenoptera^31, 70^ and other arthropods, such as Crustaceans^70, 74^. In several Hymenoptera species, clusters of CpG methylation are found across the exon-intron boundaries, as we tend to observe here^66^, which may contribute to alternative splicing via its presumed role of exon-intron “tagging”^28, 68^. Several studies in arthropods indicate a potential role of gene body methylation in transcription elongation and alternative splicing^19^, based on the apparent correlation of CpG methylation with alternative splicing found in honeybees^30, 67, 75^ and ants^69^. However, evidence from multiple insect orders suggests that CpG methylation is not directly correlated to differential exon usage^31, 58, 68^. The mixed evidence on the potential involvement of gene body methylation on alternative splicing indicates the need for future methylation studies in bumble bees that explore the possible link between CpG methylation and differential exon usage by utilizing complementary gene expression and methylation datasets.

One consistent pattern in insects is that gene body methylation is believed to be associated with unimodal expression of highly expressed “housekeeping” genes^19, 31, 59, 68, 76^. These highly expressed “housekeeping” genes are uniformly (i.e., not developmental stage- or tissue-specific) expressed^31^, exhibiting low variability in their expression pattern^68, 77^. Gene ontology analysis results from our study also support this as we noticed functional enrichment of many important essential activities in our top summarized GO term clusters, such as biological processes related to the regulation of gene expression, alternative splicing, metabolism, development, and other fundamental aspects of cell machinery. Highly methylated genes in other arthropods^26, 67–70, 77–79^ also exhibit functional level enrichment of essential cellular functions such as metabolism, mRNA processing, organelle function and transport related terms. Thus, the extent and the functional properties of gene body methylation in B*. vosnesenskii* complement the similar patterns observed in other holometabolous insects exhibiting overall low global methylation and clustered exon-biased gene body methylation, in contrast to the relatively higher global methylation and higher methylation levels extending to other genomic features (e.g., promoters, introns, and transposable elements) in hemimetabolous insects^19, 31^.

Several insect studies also suggest a link between gene body methylation and other epigenetic mechanisms^80^. For example, nucleosome dynamics, histone post-translation modifications, and associated changes in local chromatin state^81^ have been hypothesized to act in concert with CpG methylation to mediate the extent and timing of access to the transcriptional machinery and, thus, regulate subsequent gene expression levels^82^. Our data support potential cooperation among these epigenetic mechanisms as GO analysis of highly methylated CpGs included two histone modification-related terms within the top ten summarized GO terms. In insects, CpG methylation is strongly associated with histone post-translational modification and transcriptionally active chromatin marks^80, 83^. It may play a critical role in ensuring the constitutive expression of highly methylated genes across insect lineages via the exclusion of a chemically modified TSS-associated histone variant (H2A.Z) that exhibits a negative correlation to gene expression activity^28^. Thus, high CpG methylation concentration patterns of near TSS and subsequent 5′ bias could be potentially linked to CpG methylation-mediated chromatin remodeling near TSS^80^. Methylation levels in arthropods can also be related to nucleosome occupancy around the TSS, with nucleosome occupancy exhibiting positive correlations with CpG methylation^31^. No nucleosome positioning data is available for bumble bees yet; however, we hypothesize that distinct distribution pattern of distance to TSS for both highly methylated sites and differentially methylated sites observed in *B. vosnesenskii* could be potentially linked to nucleosome occupancy, especially given differences in methylation levels observed between the populations. Future multi-omics studies examining the multiple components of individual-specific epigenomes would be especially advantageous to address knowledge gaps relating to the total epigenetic landscape and regulation of context-dependent gene expression.

The second objective of this study was to evaluate the potential for differences in methylation levels among *B. vosnesenskii* from the spatial-environmental extremes of its broad distribution. We identified 2,066 differentially methylated sites between the two populations and the genomic distribution of these differentially methylated CpGs matched the trends of general CpG methylation, and were similarly overrepresented in gene bodies, especially in exons, consistent with the distribution of differentially methylated sites between sexes and castes in the bumble bee *B. terrestris*^58^. The colder high-elevation Oregon site exhibited lower percent methylation (1.03% ±0.04% SD) than warmer southern low-elevation sites in California (1.17% ± 0.06% SD), and 87.6% of differentially methylated sites were hypomethylated in the northern high-elevation samples. Although our results must be evaluated in additional populations for robust conclusions, several insect studies have reported a propensity for hypomethylation at low temperatures, including reduced CpG methylation under low-temperature stress in the tick *Haemaphysalis longicornis*^84^ and under relatively low rearing temperatures in the cockroach *Diploptera punctata* ^85^. Interestingly, while highly methylated genes are evolutionary conserved, hypomethylated genes are often faster evolving, and can be order-, genus- or species specific^68, 70^ and exhibit tissue^71^ or developmental stage specific^68^ expression. Thus, hypomethylated genes may be more plastic, exhibiting more variability and flexibility regarding their adaptability towards environmental cues^86^. The reduced methylation observed in the high-elevation Oregon *B. vosnesenskii* population is intriguing given that this population was also found to have the broadest range in critical thermal limits in laboratory experiments (CT_MIN_ vs CT_MAX_), and also exhibited the most unique gene expression patterns, especially at CT_MIN_^56^. Although we cannot link our detected methylation levels directly to thermal tolerance with the current dataset, the differences we observe between the latitude-altitude extremes in this study indicate that future studies involving CpG methylation and complementary gene expression data from specimens sampled across the altitudinal and latitudinal gradients of its wide species range would be advantageous^2^.

Genes harboring at least one differentially methylated CpG were enriched for GO terms related to several biological processes such as metabolism, reproduction, cell cycle process, and fundamental cellular activities and molecular functions including binding, transmembrane transport, and various enzymatic functions. These results are broadly consistent with gene ontology analysis of differentially methylation between reproductive states^59^ or during colony development^60^ in *B. terrestris*. Similar functional enrichment results have also been reported in differentially methylated gene sets from abiotic stress response-related studies involving silkworm^87^, water fleas^88^, and ticks^84^. Numerous GO terms overlap with previous population genomic and thermal stress studies in *B. vosnesenskii*^55, 56^, including cellular communication and functions related to responses to stress and stimuli (see Results). Of particular interest from the perspective of thermal tolerance, we observe GO terms related to response to stimulus were within the top ten summarized biological function-related clusters for both highly methylated and differentially methylated gene sets, as were terms related to calcium transporter activity (e.g., GO:1905056, GO:0005388). At the gene level, at least one differentially methylated CpG was observed in ion channel and membrane transport-related genes [*sodium/calcium exchanger regulatory protein 1-like* (LOC117234134), *TWiK family of potassium channels protein 7* (LOC117238582), *chloride channel CLIC-like protein 1* (LOC117236045), *calcium homeostasis endoplasmic reticulum protein* (LOC117242823)] and heat shock protein-related genes [*97 kDa heat shock protein* (LOC117234768), *heat shock protein 83-like* (LOC117235089)]. Heat shock protein machinery^89–91^ and ion channel/transmembrane transport mechanisms^92^, especially those linked to calcium regulation^93^ are widely recognized for their essential role in mediating molecular responses to thermal stress^93, 94^, and have been previously observed in *B. vosnesenskii*^55, 56^. The presence of chromatin-related GO terms in the top summarized GO term list is consistent with the potential involvement of CpG methylation in mediating access transcription machinery and particularly with a previously reported case of enrichment of chromatin related GO terms for differentially methylated genes related to caste determination in bumble bees^58^. Although the potential link between differential methylation and differential expression is still unclear in insects as there is mixed evidence if the differential methylation is positively correlated to differential expression^95, 96^ (but see^97–101^) or differential exon usage^59, 72^, these reported genes from our study could serve as promising candidates to more closely examine in future studies of thermal stress or other niche specific gene expression regulation in bumble bees.

Finally, our third objective was to explore the potential link between genomic and epigenetic variability in *B. vosnesenskii*. Interestingly, *B. vosnesenskii* appears to exhibit variation in thermal tolerance among populations with minimal genome-wide population structure^56^. Although we observe weak differentiation in both genome-wide SNP polymorphisms and CpG methylation, there is substantial range-wide genetic connectivity between the populations selected for WGBS (F_ST_ = 0.025). There was also no substantial correlation between methylation differences and F_ST_ in 1kb windows across the genome, especially once constitutively methylation-free regions were removed, indicating that regions with variable methylation are not located in high- or low-differentiation regions. This is consistent with a recent study in another insect, *Diploptera punctata*, which also failed to find any correlation between genetic and epigenetic variability^85^. We did observe a significant negative correlation between nucleotide diversity (π) and methylation levels across genomic windows, which is consistent with the elevated levels of methylation in gene bodies, as protein-coding regions tend to have lower levels of variation, including reduced nonsynonymous π in *B. vosnesenskii*^5^. Methylation analysis in lab-reared bumble bees also reported a potential relationship between evolutionary sequence conservation and CpG methylation^58^. While CpG methylation can potentially act as a mutagen on individual cytosines^102, 103^, paradoxically, CpG methylation in insects is enriched in evolutionary conserved “housekeeping” genes^31^ where it may play a counterintuitive role as a stabilizing factor^58^. The potential complex relationship between underlying genomic diversity and epigenetic variability in bumble bees should be further investigated, ideally including other species with more variable heterozygosity or population structure^5^.

In summary, our study provides the first high-coverage methylation profiling in a widespread North American bumble bee, *B. vosnesenskii*, and unravels the key characteristics and trends of CpG methylation in this montane species. By utilizing wild-caught specimens from the spatial-environmental species range extremes, we provide the first look at the potential for ecologically associated epigenetic variation in this native pollinator and identified multiple constitutively and variably methylated CpGs across the *B. vosnesenskii* range. *B. vosnesenskii* is a crucial pollinator and one of two species available commercially to be used for greenhouse crop pollination in North America^54^ and is also one of few North American bumble bees that may benefit from projected future climate change scenarios^51^. Although more work is needed, understanding region-specific genomic and epigenomic variation, particularly their connection to thermal adaptation, may hold considerable economic and practical conservation value. Epigenetic variation is only recently beginning to be evaluated in bumble bees, nevertheless, given the substantial colony-specific variation in bumble bee methylomes^59^, it is also possible that environmentally associated colony-specific “epi-alleles” at the population level^104^ may exist and play a role in niche-specific adaptation or may contribute to phenotypic plasticity. Further, our study only evaluated one tissue type which, while relevant for thermoregulation and flight, should be expanded to incorporate additional tissues to fully understand variation in the methylation landscape in *B. vosnesenskii.* Overall, this study provides baseline data for future studies that will include integrative multi-omics approaches (e.g., genomics, transcriptomics, epigenomics, metabolomics) from field and laboratory experiments to build a conceptual framework on the interplay between multiple modes of non-genomic epigenetic variations and its influence across multi-level molecular processes that are mediating tolerance to a broad set of environmental conditions in this species^2, 105^.

## Methods

### Samples

DNA was extracted using Qiagen DNeasy kits from the thoracic tissue of worker bees from a previous study^5^ which were collected from southern California at low elevations (CA01.2015: Latitude 36.458°, Longitude -118.879°, Elevation 314m) and from northern Oregon at high elevation (OR05.2016: Latitude 45.332°, Longitude -121.670°, Elevation 1,699m). These sites generally represent warm and cold extremes of the species range^56^ (Fig. 1a). All *B. vosnesenskii* workers (Fig. 1b) represent unique colonies based on inferences of relatedness from reduced representation SNP data^5^.

### Whole genome methylation sequencing and WGBS data analysis

Whole genome methylation libraries were prepared using the Swift AccelNGS Methyl-Seq DNA library approach for bisulfite-converted DNA (with lambda control genome spike-in) and they were sequenced on an Illumina Hiseq X sequencer by HudsonAlpha Institute for Biotechnology Genome Services Lab (Huntsville, Alabama, USA). Samples (n=8) were run in individual lanes to generate 2 x 151bp paired-end libraries. 3.6 x 10^7^ raw read pairs and 1,088Gbp in total were sequenced in the raw WGBS data set (per sample mean = 450.19 x 10^6^ ± 50.21 x 10^6^ SD read pairs and 135.96Gbp ± 15.16Gbp SD of sequence). Quality assessment of the sequenced specimens was conducted using FastQC v.0.11.9^106^. Based on the generated sequence quality assessment and a large amount of sequence data, we conducted stringent quality filtering, including adapter removal, quality trimming, removal of short sequences (< 50bp) and removed specific fixed lengths from both 5′ and 3′ ends to minimize bisulfite conversion bias using Trim Galore! v.0.6.6^107^; custom command line parameters:--illumina –q 20 – clip_R1 20 –clip_R2 20 –three_prime_clip_R1 20 –three_prime_clip_R2 60 –length 50). After stringent trimming and quality filtering of these high coverage data, we discarded ∼34.27% of raw reads, resulting in 295.90 x 10^6^ ±107.65 x 10^6^ SD trimmed read pairs and 52.33Gbp ±19.32Gbp SD per sample. We generated post-trimming sequence quality reports and sample-specific statistics using FastQC v.0.11.9^106^ and SeqKit v.0.15.0^108^. Because samples were sequenced in very high coverage and the total number of reads varied, we normalized coverage before analyses to reduce possible biases in methylation calling and subsequent analyses. Read coverage was reduced by random subsampling using SeqKit v.0.15.0^108^ to match the smallest number of read pairs in any sample (n=187,618,210 read pairs per sample). After performing the read-pair subsampling across samples, 187.62 million read pairs for each sample were aligned to the *B. vosnesenskii* genome assembly, which resulted in 75.70 ±3.04 SD fold sequencing depth per sample.

Subsampled read pairs were aligned to the *B. vosnesenskii* genome assembly (RefSeq accession GCF_011952255.1^64^) using bwa-meth v.0.2.2^109^ and alignment files were sorted using SAMtools v.1.9^110^. PCR duplicates were removed using MarkDuplicates from Picard tools v.2.23.9^111^. Methylation extraction in the CpG context from sorted post-processed BAM files was conducted using MethylDackel v.0.6.1^112^ by setting an absolute minimum coverage and employing bioinformatic removal of CpGs that were potentially C-to-T variant sites using the following parameters (--minDepth 10 –maxVariantFrac 0.5 – minOppositeDepth 10 –methylKit). Bioinformatic removal of probable C > T variants by MethylDackel resulted in the exclusion of 64,847.63 ± 5,138.26 SD CpGs per sample and resulted in a methylation call dataset containing 22,189,312.75 ± 1,919,429.19 SD CpG locations per sample. Further processing was conducted in R v.4.1.3^113^ utilizing *methylKit* v.1.20^114^. We removed CpGs with <10x coverage and with unusually high coverage (>99^th^ percentile) to minimize the effects of paralogs or repetitive regions, which excluded 1.04% ± 0.01% SD sites from the samples and resulted in 21,961,863.38 ± 1,898,407.22 SD CpGs per sample. We calculated the per base read coverage and per base methylation statistics for each sample before and after filtering using the *getCoverageStats* and *getMethylationStats* functions in *methylKit*, respectively, and utilized the average percent methylation per CpG site matrix for tabulating genome-level sample-specific and population-specific mean percent CpG methylation. There remained some dissimilarity of per base coverage within and across the samples even after read normalization, so we also normalized the coverage of the CpGs per sample using the *normalizeCoverage* function (method = “median”) in *methylKit*. We then obtained a united methylation call dataset for all samples using the *unite* function in *methylKit* that included all CpGs present in every sample at ≥10x coverage (n = 14,627,533 CpGs). As the presence of C>T SNPs can impact the accuracy of detected methylation levels in CpGs^115^, in addition to using a built-in bioinformatic detection in MethylDackel v.0.6.1, we also filtered sites using SNP data from whole genome sequencing in these populations (see the following section: “Whole genome resequencing and variant calling”). We excluded 44,041 CpGs that overlapped SNP positions so that we could focus on sites that should only be affected by methylation. After filtering, the final dataset used for general and differential methylation analysis contained 14,583,492 CpGs containing no missing data (i.e., sites that are present in every sample). Although the consistent patterns of low and similarly distributed methylation in all samples indicated successful WGBS (see Results), we repeated bioinformatic analyses by mapping reads to Escherichia phage Lambda (NCBI GenBank accession J02459.1) to assess bisulfite conversion efficiency.

To investigate the general differences in methylation among samples, we conducted principal component Analysis (PCA) using the *PCASamples* function in *methylKit* by utilizing all CpGs (n= 14,627,533) sequenced in at least 10x coverage. We also used the same CpG dataset to conduct hierarchical clustering analysis by calculating a correlation matrix from per base percent methylation data utilizing Ward’s minimum variance method with the *clusterSamples* function in *methylKit*.

### Analysis of constitutive patterns genome-wide CpG methylation in *B. vosnesenskii*

We calculated the percent methylation per CpG site (percentage reads at each CpG cytosine with a C or T) for each sample using *percMethylation* function of *methylKit*. Based on the average percent methylation for each CpG site, we categorized these sites into three categories; methylated (with ≥50% methylation); sparsely methylated (10-50% percent methylation), and unmethylated (≤10% percent methylation). We first calculated the distance from the nearest transcription start site (TSS) for all CpGs (*getAssociationWithTSS* function of *methylKit* from the NCBI *B. vosnesenskii* RefSeq annotation^64^). We used a two-sided two-sample Kolmogorov–Smirnov test to compare distributions of the distances from TSS of highly methylated sites and all CpGs using *ks.test* function in R v.4.1.3. We then used the NCBI *B. vosnesenskii* RefSeq annotation^64^ to generate feature-specific custom annotation genome tracks [i.e., Untranslated Regions of exon (exon UTR), Coding Sequences (CDS), Intron, Upstream flanking regions (Upstream Flank), Downstream flanking regions (Downstream Flank), long non-coding RNA (lncRNA), Transposable elements (TE), intergenic] following previously described methods^116^ and publicly available codes (available at: https://github.com/RobertsLab/project-gigas-oa-meth). We produced feature-specific breakdowns for all three CpG subsets (i.e., highly methylated, sparsely methylated, and unmethylated CpGs) and all sequenced CpGs. To test for statistically significant enrichment of highly methylated CpGs and the overall abundance of sequenced CpGs in each genomic annotation feature, for each feature class, we compared all CpGs against methylated sites using a Pearson’s Chi-squared test with Yates’ continuity correction implemented by *chisq.test* function in R.

After initial analyses confirmed that most methylated CpGs were confined to gene bodies, we next investigated the breakdown of CpGs based on their location within the gene body. To avoid complications that may arise from the existence of multiple transcripts due to alternative splicing, we selected the annotation of the longest isoform for each gene using the AGAT genomic toolset v.0.8.0^117^ and tabulated the fine-scale gene-body feature annotation count for each exon and intron. CpG counts for each exon and intron for protein-coding genes and long non-coding RNAs were conducted using custom bash scripts.

### Differential methylation analysis

To conduct the differential methylation analysis, we first calculated the mean and standard deviation of all CpGs using *rowSds* and *rowMeans2* function of R package *matrixStats* v.0.62^118^. Because the majority of CpGs in the genome were found to be unmethylated, as is typical for insects^27^, we removed low-variability CpGs (i.e., within less than 2 standard deviations of per base percent methylation calculated for each CpG site location across all samples) as they are not informative for differential methylation and would increase the total number of comparisons for significance testing. Overall, 93.82% of CpGs were excluded in this process. The remaining variable (SD > 2) CpGs (n=901,868) were used in differential methylation analysis in *methylKit* v.1.20^114^. We implemented Fisher’s exact test (Chi-square test) to test for significance between two population groups with basic overdispersion correction^119^ along with a false discovery rate (FDR) correction using the Benjamini-Hochberg (BH) procedure. We considered a site as differentially methylated only if there was ≥ 10% methylation change between two populations with an FDR corrected q ≤ 0.01. We defined the CpGs as “hypomethylated” and “hypermethylated” when we found statistically significant lower and higher levels of percent methylation difference in OR samples compared to CA samples, respectively. We additionally computed the number of differentially methylated sites at two other criteria (5% and 25% methylation change) to compare the pattern with CpGs identified at the 10% methylation change presented in the main Results text.

To compare the sample-specific methylation patterns, we also tabulated distances from the nearest transcription start site (TSS), compare distributions of the distances from TSS of differentially methylated sites with all CpGs, principal component analysis (PCA) and hierarchical clustering analysis for both variable (SD > 2) CpGs (n = 901,868) and differentially methylated sites (n = 2,066; assessed at 10% methylation difference) using the methods described in “Analysis of general methylation patterns” section above. We then annotated the differentially methylated sites (n = 2,066) and investigated the exon-intron breakdown of these differentially methylated sites using the methods described above and used the chi-square based contingency test as above to examine if the annotation-specific distribution of differentially methylated sites (assessed at 10% methylation difference) is significantly different than the distribution of all CpGs sequenced in the WGBS data set.

### Whole genome resequencing and variant calling

We used additional samples from the two bisulfite sequencing localities for whole genome resequencing to characterize genome-wide diversity and differentiation and identify genome positions with SNPs that could be artifactually inferred as differential methylation. We selected *B. vosnesenskii* individuals from each locality (8 for CA01.2015, 9 for OR05.2016; Fig. 1) that represent unique colonies based on inferences of relatedness from reduced representation data^5^. DNA was extracted from thoracic muscle using DNeasy kits as above and provided to the University of Oregon Genomics & Cell Characterization Core Facility for library preparation and sequencing on an Illumina HiSeq 4000 instrument. Resulting sequence data were filtered for quality using bbduk v.37.32^120^ to remove adaptors, trim low-quality bases, and remove short reads (ktrim=r k=23 mink=11 hdist=1 tpe tbo ftm=5 qtrim=rl trimq=10 minlen=25). Reads were mapped to the *B. vosnesenskii* reference genome (RefSeq Accession GCF_011952255.1)^64^ using BWA v.0.7.15-r1140^121^. SAM files were converted to the BAM using SAMtools v1.10^110^ and Picard Tools v.2.23.9^111^ was then used to sort, mark duplicates, and index BAM files. To identify a SNP set for filtering methylation data (see above) we used freebayes v.1.3.2^122^. We filtered the resulting VCF with VCFtools v.0.1.13^123^ to remove indels and non-binary SNPs, scored genotypes with depth < 4x as missing, and then retained sites with ≤ 10% missing data, Q ≥ 20, and a minor allele count of ≥ 2. Finally, we removed a SNPs with unusually high sequencing coverage (>2 times mean coverage per site) and with significant heterozygosity excess using Bonferroni correction (--hardy flag in vcftools) (following^124^). The resulting VCF included 1,162,015 SNPs after filtering, with a mean sequencing coverage of 9.97 ± 1.68 SD reads per SNP per individual and a mean missingness of 1.78% ± 1.40% per SNP per individual (98.2% complete data matrix).

The called SNP set was needed for filtering methylation calls, however genetic diversity and population structure analyses used a genotyping-free approach in the software ANGSD v.0.935-53-gf475f10^125^. ANGSD employs methods to estimate population genetic statistics from BAM files while accounting for genotype uncertainty associated with high throughput sequencing data^126, 127^. We estimated the folded site frequency spectrum (SFS)^127^ across 151 genome scaffolds of at least 100kb in length (total genome size analysed = 241,826,154bp). We estimated nucleotide diversity (π) for the two populations separately and combined using the angsd-doSfs command with minimum mapping and base quality of 20, mapping quality downgrading of C=50, GATK^128^ genotype likelihoods (GL=2), and base quality recalibration (baq=1). We then ran the realSFS program with the-fold 1 option to produce a folded SFS and thetastat –do_stat to estimate diversity parameters per site and in stepping windows of 1kb (window and step both 1,000bp). Narrow windows were used due to the rapid breakdown of linkage disequilibrium in bumble bee genomes^61^ and to avoid dilution of signal in comparisons with bisulfite data due to the globally sparse but locally clustered methylation in the *B. vosnesenskii* genome (see Results). Weighted F_ST_ was determined for the two populations by estimating the folded 2D SFS using the realSFS program and was determined per site and for 1kb windows (window and step both 1,000bp). Confidence intervals around mean nucleotide diversities and F_ST_ were obtained by nonparametric bootstrapping (1,000 replicates across windows with 1,000 sequenced sites) in the R package *boot* v.1.3-28^129^. Population structure was visualized using PCA with the PCAngsd v.1.03 program^130^ from ANGSD genotype likelihoods.

For genomic window-based analyses, we retained windows with complete sequence data across 1kb, and for comparison with methylation data, we only retained windows with at least one CpG. To test for a significant effect of methylation counts and π (log-transformed) per 1kb window, we used the R package *glmmTMB* v.1.1.5^131^ to perform a zero-inflated generalized linear model (family=negative binomial 2 to account for overdispersion). We also tested the relationship for the proportion of highly methylated (>50% category) CpGs and π (log-transformed) within each window using zero-inflated logistic regression.

### Gene Ontology (GO) analysis of highly methylated and differentially methylated CpGs

To understand the putative functional roles of genes carrying CpGs, we conducted a gene ontology (GO) analysis of two different gene sets of highly methylated and differentially methylated sites, respectively. Because there are substantial numbers of unique genes (n = 6,010) with at least one highly methylated CpG site represented in the gene set, we decided to set a predefined criterion (i.e., use the subset of unique genes harboring a minimum of 100 highly methylated sites) to conduct functional enrichment analysis. Based on this criterion, we selected a subset of unique genes (n=44) which were subsequently used in our gene ontology analysis. We also conducted a separate functional enrichment analysis where we included all unique genes (n = 1,272) harboring all differentially methylated sites (n = 2,066) assessed at 10% methylation difference. We conducted functional enrichment analysis for both gene sets using R package *GofuncR* v.1.14^132^ and utilized the curated B. *vosnesenskii* GO annotations from Hymenoptera Genome Database^133^. We considered the GO terms significant at P < 0.05. We used semantic similarity-based reduction of GO terms and visualized the enriched GO term list using GO-Figure!^134^. We independently compared the statistically significant GO terms, and representative summarized GO terms (i.e., clusters) from both gene sets with GO term lists from two previous studies^55, 56^. In one of these studies, Pimsler et al.^56^ identified 1786 enriched statistically significant GO terms (assessed at P ≤ 0.05) for seven different contrasts and directions of gene expressions). We combined these GO term lists into a single list representing the unique GO terms (n=1398) found at least once in any of these contrasts to compare them with our study’s two individual GO term lists. We also compared our gene ontology (GO) results with another study^55^ by Jackson et al. which provided two different enriched GO term lists from outlier gene lists detected from tested for associations with variable temperature (n= 151 GO terms) and precipitation (n=86 GO terms). We combined these two GO term lists into a single list representing 221 unique GO terms from both categories and compared them with GO term lists from our study.

## Acknowledgements

This work was supported by National Science Foundation for under grants no. DEB 1457645 and URoL 1921585 to awarded to J.D.L. Any opinions, findings, and conclusions or recommendations expressed in this material are those of the authors and do not necessarily reflect the views of the National Science Foundation. We thank Jason M. Jackson for his assistance with specimen collection and sample processing. We also thank Lozier lab members for useful their helpful feedback and discussion.

## Author contributions

JDL obtained funding, performed WGS bioinformatic and statistical analysis, and supervised the project. SRR designed and conducted bioinformatic and statistical analyses for WGBS and functional genomic approaches. SRR wrote the manuscript with contributions from JDL. Both authors have contributed, read, and approved the final manuscript.

## Data availability statement

Raw WGBS reads generated in this study has been deposited and is currently available at the National Center for Biotechnology Information (NCBI) Sequence Read Archive (SRA) under NCBI BioProject PRJNA956115.

